# Comprehensive protease specificity profiling

**DOI:** 10.1101/2024.11.06.622033

**Authors:** Bo Zhu, Michael D. Lane, Joshua A. Baller, Pratik D. Jagtap, Timothy J. Griffin, Burckhard Seelig

## Abstract

Protease enzymes are of great importance in medicine, industry, and as research tools. Despite the crucial need for detailed knowledge of their proteolytic cleavage specificity, many proteases are poorly characterized. We present a method for fully characterizing the cleavage specificity of proteases through the comprehensive profiling of all possible permutations of octamer peptide substrates in a single experiment. The powerful combination of in vitro selection with high-throughput sequencing, mass spectrometry, and automated motif mining enabled the screening of mixtures of around 10^12^ peptides. We developed freely available software that easily integrates the massive amounts of cleavage data into user-friendly specificity information. We applied this method to three different proteases that had either hard-to-study broad specificity (ADAM17 and streptopain) or narrow specificity (factor Xa) as a control. The resulting specificity maps revealed motifs that corroborate canonical known cleavage sites, yet step further into extended spectrum preferences and yield insights into the function of broad specificity proteases.

## Introduction

Proteases perform a vast array of critical functions in the cell. These activities are dependent upon the highly-refined specificity of the protease, including evolved natural cleavage specificities and co-evolved cleavage sites ^1^. Recent efforts have assessed broad-specificity proteases using human proteomic screens ^2-11^, designed peptide libraries ^12^, or photo-crosslinked native substrates to protease ^13^. However, these methods have drawbacks that limit their ability to generate reliable specificity maps for broad specificity proteases, including low numbers of tested peptides (usually dozens, up to thousands), proteomic library bias, or the need for modified or unnatural amino acids in the cleaved substrates. To increase the throughput of protease cleavage site analyses, in vitro selection by phage display was also used ^14-17^. However, even the largest phage-displayed peptide library reported still can only test 4% of all possible octamer substrate sequences and the potential cleavage of the g3p protein that is crucial for displaying the substrates on the phage introduces bias. Furthermore, the reported method requires prior knowledge of the protease specificity to perform the cleavage site-guided motif generation ^16^. Here we present a method that overcomes these limitations by combining a complete *in vitro* selection by mRNA display, high-throughput sequencing, and LC-MS/MS to increase the number of analyzed cleavage sites by several orders of magnitude, eliminate proteomic or modified amino acid biases, provide knowledge-free cleavage site-guided alignment, and enable screening of all possible substrate octamers in a single experiment without repeated selection rounds (Fig. 1).

**Figure 1.**
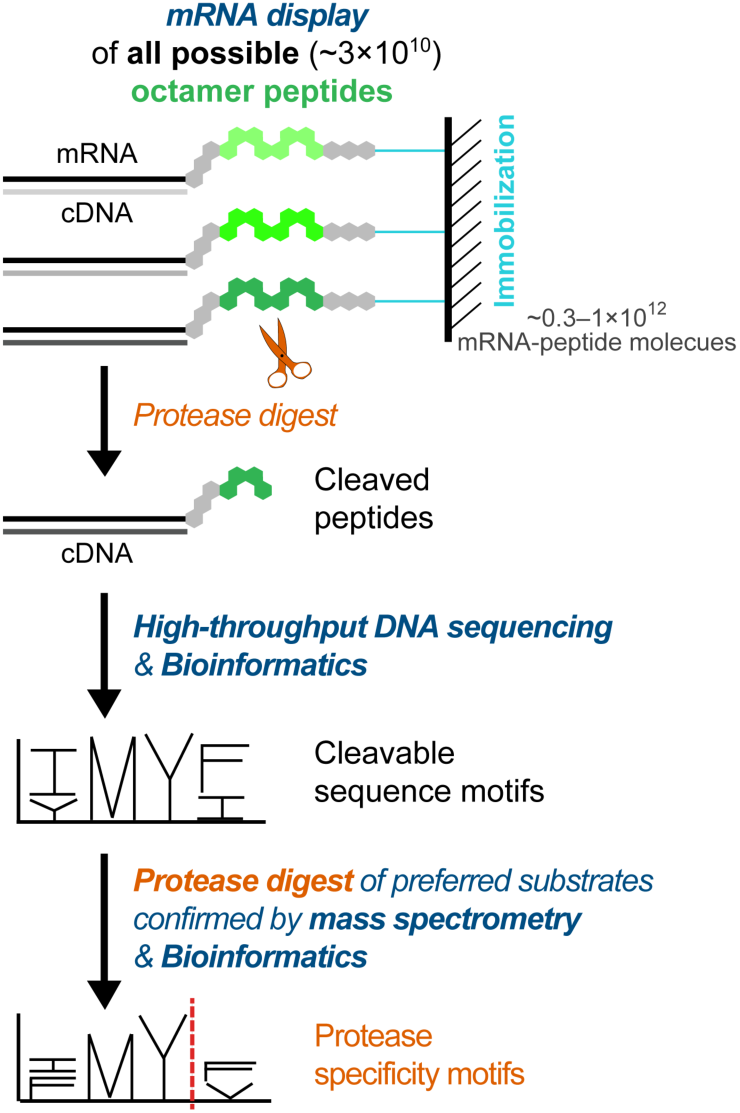
General scheme for comprehensive protease specificity profiling by mRNA display, high-throughput sequencing, LC-MS/MS, and automated motif mining. A library containing all octamer peptide substrates is mRNA displayed, immobilized, and subsequently digested with a protease of interest. The cDNA-peptide fusions released from immobilization by protease cleavage of the respective peptide substrate are analyzed by high-throughput sequencing to identify the cleavable sequences. The cleavable sequence motifs are obtained through the motif mining computation pipeline. Highly enriched sequences from the cleavable sequence motifs are chemically synthesized and the cleavage position is identified by LC-MS/MS. These cleavable sequence data and the identified cleavage positions are merged to generate specificity motifs with cleavage positions. The red dashed line indicates the cleavage position.

Proteases typically recognize between 2-4 residues on either side of the cleaved bond and, therefore, their cleavage preferences can be challenging to elucidate ^18^. To scan all possible permutations of eight residues (octamers), we used the mRNA display technology ^19, 20^, which had previously been used for substrate preference studies of post-translational modification enzymes ^5, 6, 21-23^. We generated up to 10^12^ substrates and thereby oversampled all possible octamer substrates (20^8^ = 26 billion) by > 30-fold. We applied this method to comprehensively assess the protease specificity C- and N-terminal to the cleaved bond. Subsequent high-throughput sequencing enabled us to detect up to millions of cleavage sequences for motif mining. The number of sequences is a substantial increase over the best existing method and provides high-quality big data for machine learning model building for protease studies. Finally, mass spectrometry (MS) validated the subsets of preferred substrates and identified the location of cleavage for a guided alignment of all cleaved sequences to generate the cleavage specificity motifs. Our bioinformatic tool realized the similarity-based hierarchical motif assembly, which is critical to also find minor motifs for proteases with broad specificity ^24^.

The key advantages of our method, in addition to the unique ability to sample all possible octamer substrates, are: (i) a substantial reduction of biases because our substrate library contained an even distribution of only natural amino acids and required no additional protease processing prior to selection; (ii) the simultaneous assessment of both prime and non-prime cleavage specificity; and (iii) the ability of our purely *in* vitro technology to freely assess protease function with respect to temperature, pH, ionic strength, or inhibitors ^25, 26^. Moreover, while lower-throughput methods might be sufficient to characterize narrow-specificity proteases, higher-throughput is needed to reveal the intricate details of broad-specificity proteases.

We applied this method to study a representative group of narrow- and broad-specificity, well-and poorly-characterized protease enzymes: factor Xa, ADAM17, and streptopain. The comprehensive high-resolution specificity maps generated with our method will lead to a better understanding of proteases in general, and broad-specificity proteases in particular.

## Results

### Design of the comprehensive octamer peptide substrate library

We aimed to present all possible octamer substrates for cleavage. Through mRNA display, we generated the random octamer peptide library, where every individual peptide variant was linked at one end to its encoding mRNA, and at the other end to an immobilization tag (Fig. 2a). mRNA display also requires additional constant regions on each terminus of the peptide sequence that is to be displayed ^27^. We incorporated an alkyne moiety at the conserved N-terminus to allow bioorthogonal immobilization via copper-catalyzed click chemistry ^28^. This covalent immobilization to azide-agarose resin enabled stringent washes to remove nonspecific binders during the subsequent selection step. We used an orthogonal tRNA/aminoacyl-tRNA synthetase pair to incorporate the alkyne group specifically at an amber stop codon ^29^. The entire translated peptide construct consisted of MG, alkyne moiety (Z), 6 × His (H) affinity tag, PQP frame around the random octamer substrate (X_8_), and ended with MPTGY for mRNA display purposes (Fig. 2b). PQP was chosen as a frame because this motif is generally difficult for most proteases to cleave. We constructed the random octamer section of the library (Table S1) through chemical DNA synthesis using a mixture of nucleotide trimers encoding all 20 amino acids ^30^. This strategy eliminated the risk of introducing stop codons and degeneracy bias, which can be an issue when using the more common NNS or NNK libraries. Through *in vitro* translation of this library, we were able to produce up to 10^12^ peptides as mRNA-displayed fusions, which oversamples the library of all possible octamers (2.6 × 10^10^) by >30-fold. This comprehensive library of all octamer peptide substrates can be directly used for profiling the cleavage specificity of essentially any protease.

**Figure 2.**
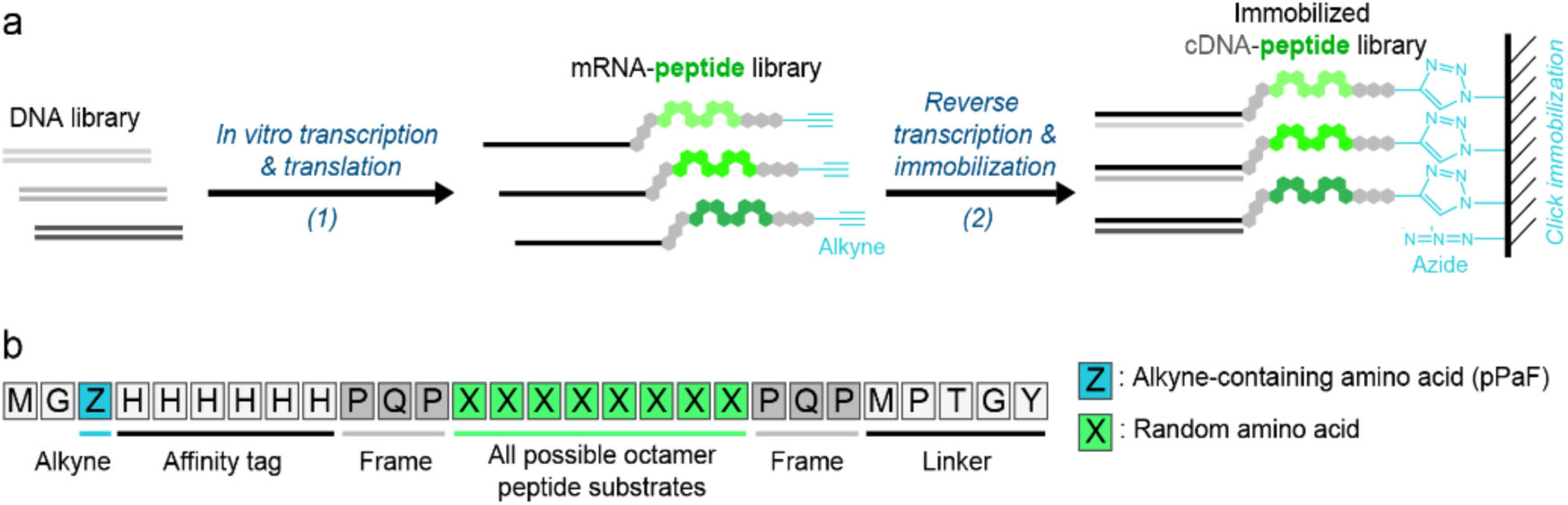
Overview of mRNA display and click immobilization of octamer peptide substrates for exploring protease specificity. (a) The strategy used for mRNA display and immobilization. (1) DNA library is transcribed to mRNA library, modified with puromycin, and in vitro translated by the PURE system to display each peptide as covalent fusion with its own encoding mRNA. The translated peptide is modified with an alkyne during translation via incorporation of the alkyne-containing non-natural amino acid pPaF. (2) The mRNA-peptide library is reverse transcribed, and the cDNA-peptide library is immobilized on agarose beads through Cu(I)-catalyzed alkyne-azide cycloaddition (CuAAC) click reaction. (b) Design of library containing all possible octamer peptide substrates with termini compatible for mRNA display.

### Isolation of cleavable peptide substrates for three proteases by mRNA display

We chose three proteases comprising both narrow- and broad-specificity enzymes to validate our new method: factor Xa, ADAM17, and streptopain. Factor Xa is a narrow-specificity serine protease and a critical component of the coagulation cascade. The cleavage specificity of factor Xa has been well characterized by several methods ^31-35^. ADAM17 is a broad-specificity transmembrane metalloprotease ^36, 37^ that is involved in development, healthy physiology, and pathological mechanisms. Known targets of ADAM17 include Tumor Necrosis Factor alpha (TNFα) and substrates involved in tumor immune-surveillance, inflammation, and cancer, implicating the protease in diseases from rheumatoid arthritis and inflammatory bowel disease to large diffuse non-Hodgkin B-cell lymphoma ^38, 39^. The specificity of ADAM17 has been characterized by a peptide library approach ^24^, Q-PICS ^40^ and SPD-NGS ^16^. Finally, streptopain is a broad-specificity cysteine protease produced by *S. pyogenes*. Streptopain, also known as streptococcal pyrogenic exotoxin B (SpeB), has potent immunomodulating effects on the human immune system, cleaving multiple immunoglobulins, critical components of both the classic and alternative pathways of complement activation, and numerous chemokines ^41^. Further, streptopain can cleave a variety of host proteins unrelated to the immune system, such as fibrin/fibrinogen, plasmin, and host matrix metalloproteases, as well as a wide range of bacterial proteins ^41^. Despite the importance of streptopain, its cleavage specificity has only been poorly characterized to date. Only about 25 quenched fluorescent peptide substrates have been used to probe positions P3-P1 and P1′, which resulted in a cleavage specificity map ^42^ that is entirely inadequate to explain streptopain’s broad activity. The limited information on streptopain in the MEROPS database ^18^ (28 substrates) is insufficient to reliably predict its behavior even towards known cleavable substrates ^42^.

We synthesized and immobilized a complete library of mRNA-displayed octamer substrate peptides to agarose beads via click reaction for each protease experiment (Fig. 2a) and washed the immobilized library with the respective digestion buffer. The subsequent incubation with protease cleaved only susceptible octamers and thereby released the cDNA/mRNA-fusions that encoded for those cleavable peptide substrates (Table S2). The released cDNA was subjected to high-throughput sequencing to analyze the region that encoded the cleavable protease substrates (Table S3). In addition, we also prepared a control sample of the library of immobilized sequences, identical to the sequences immediately before incubation with protease. This sample enabled us to assess any potential bias that might occur during translation, transcription, or immobilization (Table S1).

### Analysis of high-throughput sequencing data to identify cleavable sequence motifs

The flowchart in Fig. 3 shows the entire process of the protease specificity motif computation. Using this bioinformatics pipeline, we first analyzed the high-throughput DNA sequencing data to create cleavable sequence motifs (Fig. 3a-g), and then incorporated mass spectrometry data to yield the final protease specificity motifs, including the cleavage position (Fig. 3h-j). Filtering, trimming, and translating the high-throughput sequencing raw data yielded 13-16 million octamer peptide sequences that conformed to the library design (Fig. 3b, Table S3). To estimate the false positive rate of the detection method, a control dataset was generated *in silico* by shuffling the amino acid order of the sequences in the selected pool (Fig. 3c). Next, the k-mers (subsequences of length k) frequencies of the datasets were enumerated. A k-mer length of six was selected after observing that when longer lengths were used, specific k-mers showed up too infrequently to reliably evaluate k-mer presence and absence (Fig. 3d). The processed 6-mer sequences of the cleaved peptides were compared to the library of peptides before cleavage (control) to calculate an enrichment score for each 6-mer sequence. The 6-mer sequences below the frequency and enrichment cutoffs were filtered out (see methods section for details). A network of the surviving 6-mer sequences was generated. Sequences were pruned from the network if their number and strength of connections to others were below the specified cutoffs. This pruning was performed to consistently obtain large complex networks with the true datasets, while yielding sparse or no networks with randomized datasets (Fig. 3e). Consensus motifs were then generated by agglomerative hierarchical clustering of remaining 6-mers (Fig. 3f). The motifs were filtered by the fraction of total unique sequences they contained and overall similarity of last merge. The resulting parent motifs in each branch represented our cleavable sequence motifs (Fig. 3g, Fig. S2). These cleavable sequence motifs are shown for each of the three proteases in Figs. 4a, 5a, and 6a. The positions of those motifs within the hierarchical tree are listed in Table S4.

**Figure 3.**
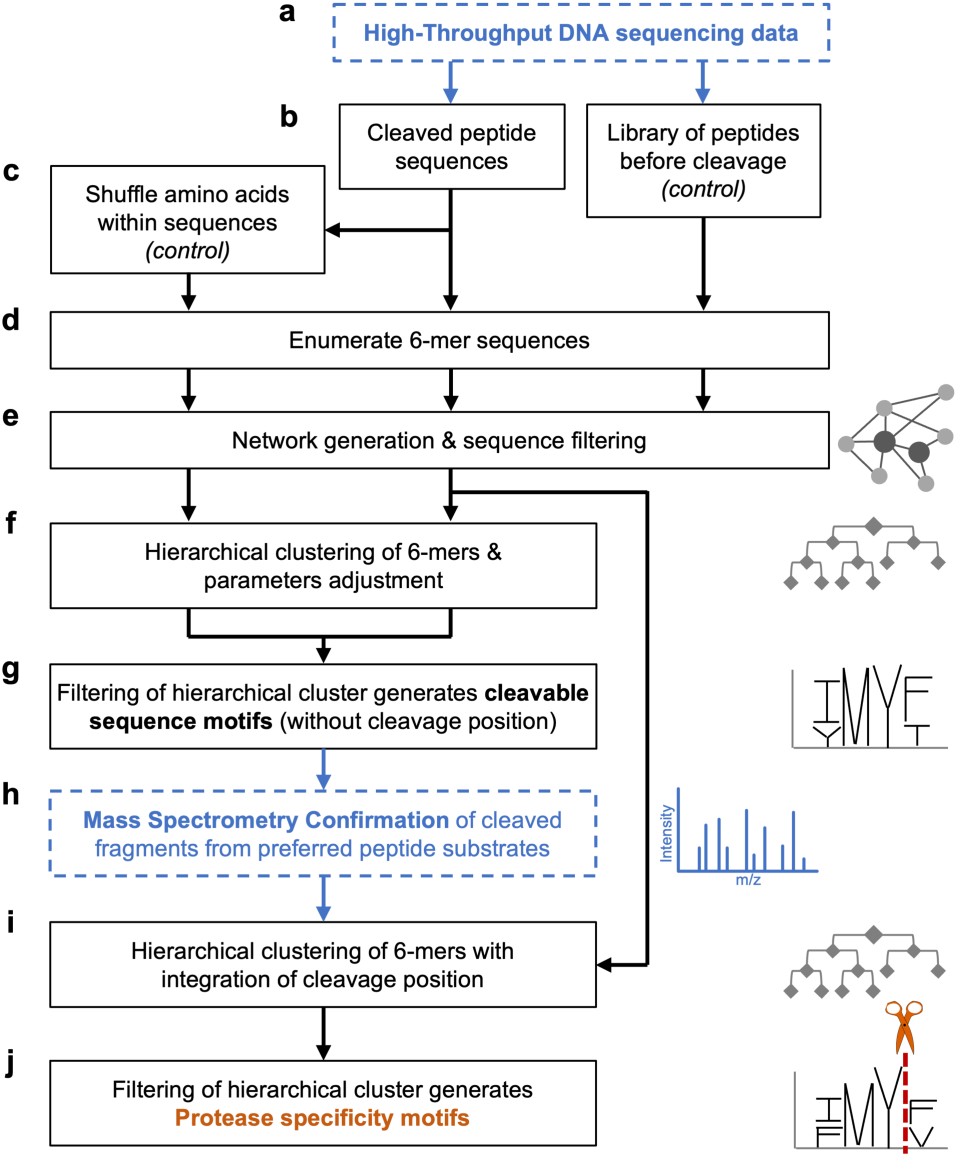
Computational motif mining pipeline. Automated processing of the high-throughput DNA sequencing data from the peptide substrate library before and after protease cleavage, and the subsequent incorporation of mass spectrometry cleavage data for preferred peptide substrates ultimately yields the protease specificity motifs for a given protease. The two data entry points are highlighted by dashed blue boxes. A flowchart with all steps in full detail can be found in Fig. S1.

**Figure 4.**
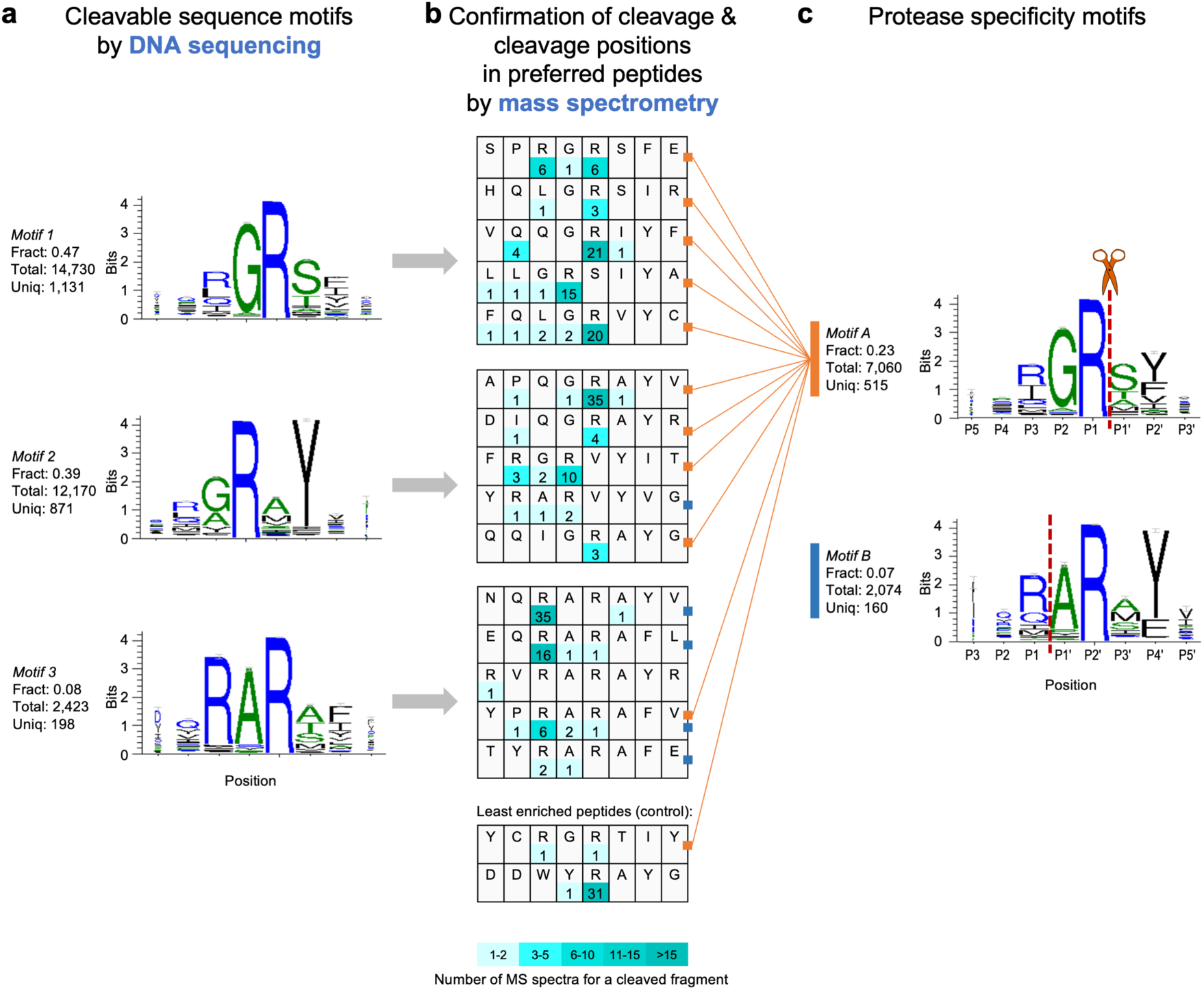
Protease specificity motif mining for factor Xa. **(a)** DNA sequencing data of cleavable peptides yielded the three most enriched cleavable sequence motifs. Total: Number of sequences supporting this motif. Fract: Sequences supporting this motif as fraction of all sequences enriched for this protease. Uniq: Number of unique 6-mer sequences supporting this motif. **(b)** Cleavage positions in preferred peptides identified by mass spectrometry (MS). Numbers represent the number of MS spectra obtained (in shades of teal). Grey arrows indicate from which motif the group of preferred peptides originated. **(c)** Protease specificity motifs with cleavage position generated by combining cleavable sequence motifs seen in (a) with MS results seen in (b). Red dashed line indicates cleavage position. Only the six highest-ranking motifs are shown. Color match between squares after peptide sequence and vertical bar to the left of a motif indicates that the motif is supported by cleavage information from this sequence (connected by orange lines for motif A as example). Motifs are ordered from top to bottom by number of unique sequences supporting a motif. Sequence logos are generated from the aligned sequences using WebLogo 3.5.0: Overall height of each stack indicates sequence conservation at that position with height of an amino acid within the stack reflecting relative frequency. Error bars are the 95% confidence interval for the reported conservation. Width of amino acid letters corresponds to the relative frequency of ‘valid’ symbols at that position (the degree of support for amino acid frequencies at that position). The color of the amino acids represents their hydrophobicity (blue - hydrophilic, green – neutral, black – hydrophobic).

**Figure 5.**
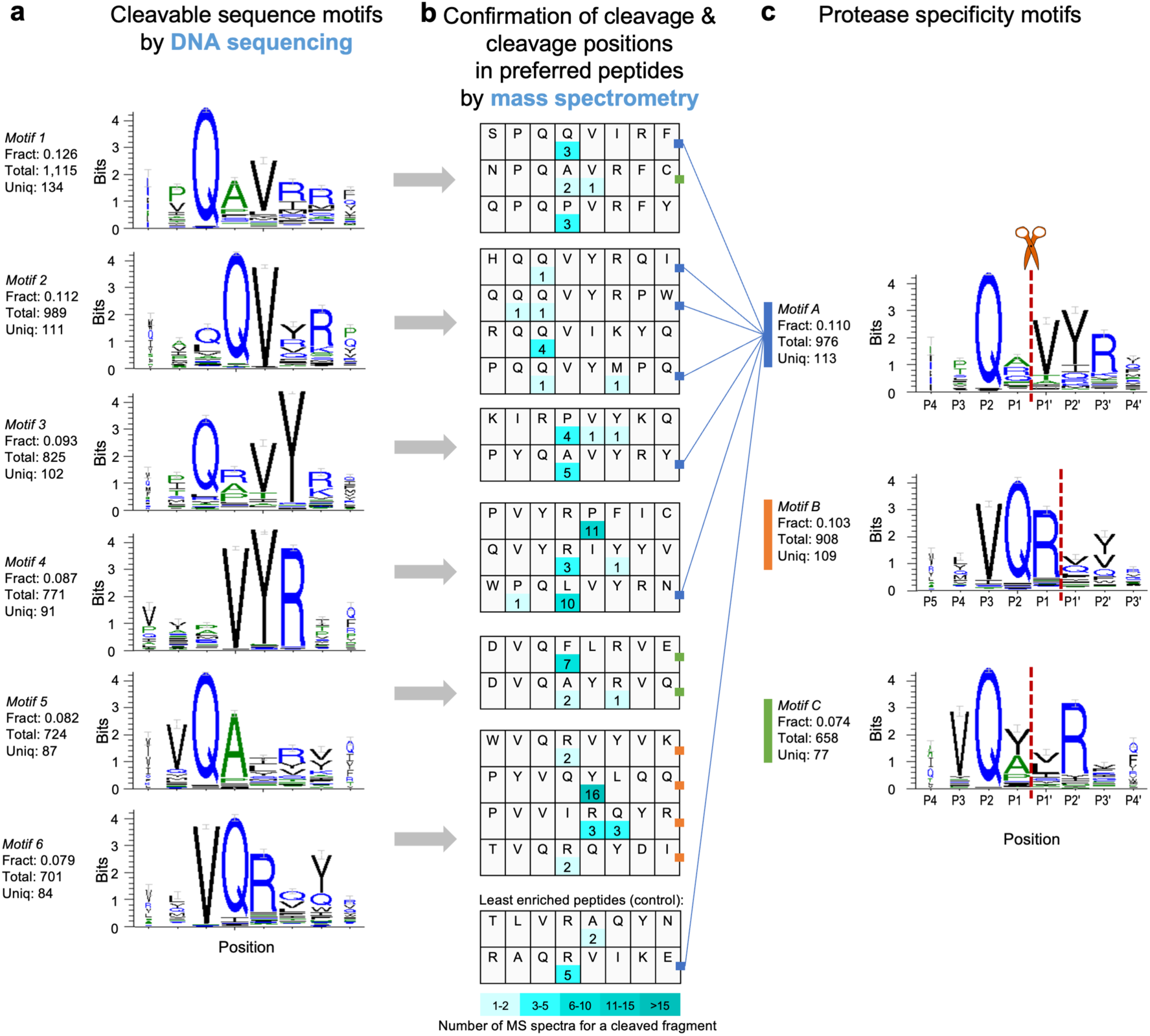
Protease specificity motif mining for ADAM17. (**a**) DNA sequencing data of cleavable peptides yielded the six most enriched cleavable sequence motifs. Total: Number of sequences supporting this motif. Fract: Sequences supporting this motif as fraction of all sequences enriched for this protease. Uniq: Number of unique 6-mer sequences supporting this motif. (**b**) Cleavage positions in preferred peptides identified by mass spectrometry (MS). Numbers represent the number of MS spectra obtained (in shades of teal). Grey arrows indicate from which motif the group of preferred peptides originated. (**c**) Protease specificity motifs with cleavage position generated by combining cleavable sequence motifs seen in (a) with MS results seen in (b). Red dashed line indicates cleavage position. Only the four highest-ranking motifs are shown. Color match between squares after peptide sequence and vertical bar to the left of a motif indicates that the motif is supported by cleavage information from this sequence (connected by blue lines for motif A as example). Motifs are ordered from top to bottom by number of unique sequences supporting a motif. Sequence logos are generated from the aligned sequences using WebLogo 3.5.0: Overall height of each stack indicates sequence conservation at that position with height of an amino acid within the stack reflecting relative frequency. Error bars are the 95% confidence interval for the reported conservation. Width of amino acid letters corresponds to the relative frequency of ‘valid’ symbols at that position (the degree of support for amino acid frequencies at that position). The color of the amino acids represents their hydrophobicity (blue - hydrophilic, green – neutral, black – hydrophobic).

### Confirmation of protease cleavage and determination of cleavage positions by mass spectrometry for motif refinement

To independently confirm that the sequences supporting the motifs are cleaved by the proteases, and to refine the motifs generated from the first stage of automated motif mining (Fig. 3g), we determined the cleavage position for a subset of peptides by a cleavage assay. For this purpose, we chose the five most highly enriched 6-mer sequences from each of the identified cleavable sequence motifs (Fig. 3g). In addition, we also chose, out of all 6-mer sequences, the five most highly enriched and the two least enriched sequences as controls (Fig. 3e). For each of these 6-mers, we extracted the associated octamer sequences from the un-processed dataset. A four amino acid long tag was added to the N-terminus of the octamers to facilitate subsequent mass spectrometric analysis by increasing charge and hydrophilicity. The peptides were chemically synthesized and combined into groups of 20 or fewer peptides each. We digested these mixtures with the respective protease. The resulting cleaved fragments were analyzed by LC-MS/MS to identify the position of cleavage for this subset of peptides (Fig. 3h; Figs. 4b, 5b, 6b). We also confirmed the cleavage position of a known substrate for each protease by adding that substrate to the digest as positive control. The mass spectrometry results revealed that most of the peptides tested had multiple cleavage sites, which is reasonable since a peptide with multiple cleavage sites is an even better protease substrate than a peptide with only a single site.

**Figure 6.**
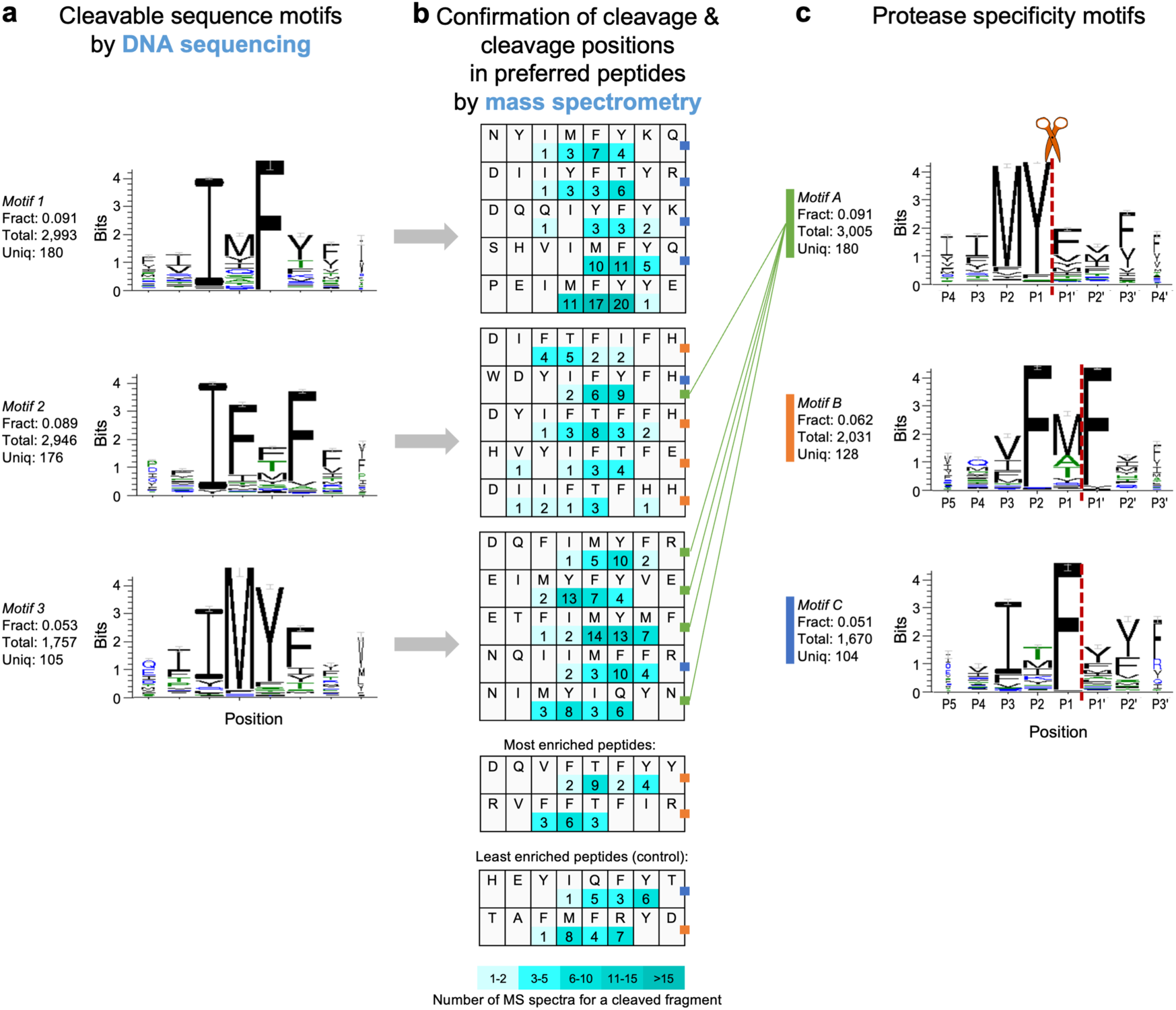
Protease specificity motif mining for streptopain. (**a**) DNA sequencing data of cleavable peptides yielded the three most enriched cleavable sequence motifs. Total: Number of sequences supporting this motif. Fract: Sequences supporting this motif as fraction of all sequences enriched for this protease. Uniq: Number of unique 6-mer sequences supporting this motif. (**b**) Cleavage positions in preferred peptides identified by mass spectrometry (MS). Numbers represent the number of MS spectra obtained (in shades of teal). Grey arrows indicate from which motif the group of preferred peptides originated. (**c**) Protease specificity motifs with cleavage position generated by combining cleavable sequence motifs seen in (a) with MS results seen in (b). Red dashed line indicates cleavage position. Only the four highest-ranking motifs are shown. Color match between squares after peptide sequence and vertical bar to the left of a motif indicates that the motif is supported by cleavage information from this sequence (connected by a green line for motif B as example). Motifs are ordered from top to bottom by number of unique sequences supporting a motif. Sequence logos are generated from the aligned sequences using WebLogo 3.5.0: Overall height of each stack indicates sequence conservation at that position with height of an amino acid within the stack reflecting relative frequency. Error bars are the 95% confidence interval for the reported conservation. Width of amino acid letters corresponds to the relative frequency of ‘valid’ symbols at that position (the degree of support for amino acid frequencies at that position). The color of the amino acids represents their hydrophobicity (blue - hydrophilic, green – neutral, black – hydrophobic).

The cleavage sites information was fed back into the software pipeline (Fig. 3h) to refine the cleavable sequence motifs and generate the final protease specificity motifs. For that purpose, the step (f) was repeated with 6-mer sequences that now included the position of cleavage (Fig. 3i). The motifs in the hierarchical cluster were filtered by the fraction of total nodes and overall similarity of last merge, and only the parent motif in a branch was kept. The filtering by similarity used in Fig. 3g was adjusted to ensure that at least one tested peptide with a known cleavage position was supporting the protease specificity motifs obtained (Fig. 3j). The three proteases factor Xa, ADAM17 and streptopain were subjected to this method. We successfully assigned the cleavage positions and obtained high-resolution cleavage specificity profiles for each protease as shown in panel c of Figs. 4, 5 and 6.

### Protease specificity motifs of factor Xa

The narrow specificity serine protease factor Xa was used as a proof-of-principle application that successfully verified the accuracy of our method against this very well-studied protease. The comprehensive results for factor Xa are shown in Fig. 4. The top three enriched cleavable motifs (Fig. 4a) were supported by 2,200 unique sequences and show the well-known GR or AR pattern ^17, 32^. Mass spectrometry of the digested preferred peptide substrates clearly confirmed the known preferential cleavage after the R residue (Fig. 4b). Even the two peptides with the lowest copy number among the enriched peptides (least enriched peptides) were digested, which confirmed that the enrichment of cleavable sequences was successful. Fig. 4c shows the factor Xa protease specificity motifs including the cleavage site. The top two specificity motifs were supported by 675 unique sequences (Fig. 4c). The top motif A alone was supported by 515 sequences. In contrast, less than 86 sequences were used previously to generate the specificity motifs for factor Xa by the PICS assay ^32^. We speculate that motif supported by the highest number of unique sequences represents the major recognition pattern, while the other motifs can be used for the discussion and exploration of subsite activities. Further, while the canonical specificity of R in P1 position and G or A in P2 were identified by PICS, it was unable to clearly conclude the specificity in positions P3 and P2′, based on the results from the two different peptide substrate libraries. Interestingly, most highly enriched and reactive sequences PQGR, IQGR, QQGR from a previously published tetramer library using phage display ^17^ were also confirmed in our list of highly enriched sequences (Fig. 4b).

The major protease specificity motif A (Fig. 4c) contained the well-characterized GR or AR pattern ^17, 32^, while motif B contained the QR pattern described in the MEROPS database. Motif A confirmed the canonical specificity for R at position P1, and for the small amino acids G and A at position P2. The specificity for P3 were R, I, Q and L, among which the R in P3 agreed with four previous studies ^17, 33, 35, 43^, and the L and Q in P3 agreed with two of them ^17, 43^. However, similar to the result from PICS, the cleavage at K at P1 described by a previous study ^18^ (collected by MEROPS motif) was not confirmed.

A unique and previously unreported Q-R (P2-P1) pattern was identified in motif B. The Q in P2 position, which we identified, was not found in the MEROPS database ^18^ nor in the motifs from the PICS method. The cleavage after QR was confirmed by mass spectrometry (Fig. 4b), and the QR pattern had also been found previously in the LLQR sequence that was highly enriched by phage display ^17^.

Few previous studies focused on the specificity of prime side for factor Xa. The motif A shows the P1′ preference for S, T and A. The S and T preference had been described in the MEROPS database (based on 63 cleavages) ^18^, while S and A had been found by PICS. Furthermore, motif A clearly demonstrates a specificity at P2′ for Y, F, V and I. In summary, the motifs obtained by our method were supported by more than 7-fold sequences compared to the motifs generated by the PICS method, or those described in the MEROPS database ^18^.

### Protease specificity motifs of ADAM17

The broad specificity ADAM17 protease was investigated here to apply our method towards a protease that has been well-studied by the most powerful previous methods for understanding broad-specificity proteases. Our comprehensive results for ADAM17 are shown in Fig. 5. The top six enriched cleavable motifs (Fig. 5a) were supported by 609 unique sequences with the QAV or QRV patterns dominating. The mass spectrometry results of the digested preferred peptide substrates showed a trend of cleavage after the residue following the Q, or RP combination (Fig. 5b). The digestion was confirmed for even the least enriched peptides, which verified again the successful enrichment of cleavable sequences. Fig. 5c showed the ADAM17 protease specificity motifs including the cleavage site. The top three specificity motifs were supported by 299 unique sequences and are shown in Fig. 5c. For comparison, the PICS assay had to rely on 256 sequences in total to generate all ADAM17 specificity motifs ^40^.

The motifs A–C showed some similarity with subsets of amino acid preferences at different positions. They all showed a specificity for Q at P2 position, which agreed with the MEROPS database ^18^, but the PICS method did not identify this preference. The amino acids A and R were preferred in the P1 position for the three motifs, which are also the top two amino acids at P1 position in MEROPS database ^18^. The V preference in the P1′ position in the three motifs is the same as found by PICS and in the MEROPS database ^18^; the L and I preference in P1′ in motif C also agreed with PICS results and one other previous study ^24^. The canonical substrate of the ADAM17 protease, TNFα (PLAQA↓VRSSS, with ↓ indicating the cleavage position) ^44^, was also corroborated by our motif A and motif C for the four positions P2-P2′. However, the PICS results only covered the three positions (P1-P2′). ADAM17 has been reported to cleave the human ACE2 (SARS-CoV-2 receptor) after R708 (RMSR↓SRIN), which we confirmed in our motif B in P1 position ^45^. The strong preference for VQR in positions P3-P1 of our motif B agrees with another native substrate of ADAM17, Cadherin-3 (EVQR↓LTVT), that was suggested by a recent secretomes analysis ^46^ but was not found by the PICS method. The VQA pattern in our motif C agrees well with a cleavable site in the EPHA2 receptor (QVQA↓LTQE) found in that same analysis ^46^. The human IgG Fc receptor III (CD16) is another well-characterized substrate of ADAM17, which has previously been shown to be cleaved three times *in vivo* (ITQGLA↓V↓ST↓ISSF) ^47^. We also observed multiple cleavages for several of our tested peptides (Fig. 5b). When CD16 is cleaved between A and V, the P1 and P1′ positions agree with our motifs A and C. When the cleavage happens after V, the P1 position agrees with two peptide digestion results in Fig. 5b, however the previously reported SPD-NGS method did not show the preference of V at P1 position ^16^. Our motif A also had Q preferred in the P1 position, which is found in another native substrate, the Interleukin 6 receptor (IL-6R) (SLPVQ↓DSSSV) ^48^. In contrast, the PICS results did not agree with this natural substrate sequence. The Ebola virus surface glycoprotein GP is also a native ADAM17 substrate (KTLPD↓QGDND) ^49^, and the Q in P1′ position agreed with our motif B, but again the PICS method did not identify a match at any position. The SPD-NGS method did not show a high confidence score for cleavage prediction for Q at either P1 or P1′ positions ^16^.

A crucial role of ADAM17 is the proteolytic processing of tumor necrosis factor alpha (TNFα), which is important in the regulation of immune cells ^38^. Interestingly, the exact P2-P2′ sequence of TNFα (QAVR) was found in one of our highly enriched 6-mer sequences: PQAVRF. To probe the human proteome for additional matches with our enriched 6-mer sequences, we searched the human proteome database using the peptide search tool UniProt ^50^ for a small subset of our >1,000 enriched sequences, namely the 20 enriched sequences with MS-confirmed cleavage position (Fig. 5b). The entire 6-mers, and their 5-mer subsequences that retained the cleavage site were used for the searches, returning 8 and 529 matches, respectively. These matches included 35 cell surface transmembrane proteins. ADAM17 is known to cleave transmembrane-anchored proteins proximal to the membrane ^51^; therefore, we further analyzed the 6-mer and 5-mer matches in known transmembrane proteins for their potential protease accessibility (defined as being located within 30 amino acids proximal to the membrane). The sequence LVRA↓Q is located in the putative olfactory receptor 52L2 (UniProt ID: Q8NGH6), which is responsible for the detection of odorants in olfactory receptor neurons ^52^. Another sequence PQQ↓VI is found within the facilitated glucose transporter member 2 (UniProt ID: P11168), which facilitates glucose movement across cell membranes ^53^. These cleavage sites are located on the protein surface according to their predicted structures by AlphaFold ^54, 55^ and should therefore be protease-accessible. These examples indicate that the highly enriched cleavable sequences identified by our method and their matches to the human proteome could help inform future research on the physiological function of ADAM17. Expanding this search for proteome matches beyond these 20 enriched peptide sequences will likely yield additional hits of potentially interesting ADAM17 substrate candidates for further studies. These likely substrate candidates will have to be confirmed by testing the respective full-length proteins for cleavage under physiologically relevant conditions. Such an analysis is beyond the scope of the current study, not least because of added challenges due to many of the candidates being membrane proteins.

### Protease specificity motifs of streptopain

The broad specificity cysteine protease streptopain was assessed by our method to further understand a protease with important clinical applications whose specificity has been notoriously difficult to analyze. Fig. 6 shows the comprehensive results for streptopain. The top three enriched cleavable motifs (Fig. 6a) were supported by 461 unique sequences, and the hydrophobic amino acids were enriched in these motifs. The mass spectrometry results of digested preferred peptide substrates revealed that multiple cleavages (>3 for most sequences) occurred in the enriched sequences (Fig. 6b). The successful cleavage of the two least enriched peptides confirmed the effective enrichment of all cleavable sequences. Fig. 6c showed the streptopain protease specificity motifs including the cleavage site. The top three specificity motifs were supported by 412 unique sequences in total (Fig. 6c). A previous study showed that hydrophobic residues are preferred at P2, but little preference was reported for positions P3 or P1′ ^42^. Another analysis by PICS also showed preference for hydrophobic residues at P2, but inconsistent results were obtained for other positions (e.g., P1 and P1’) from different proteome substrates in that study ^10^. Our motifs preferred M/F/V/Y (motif A), or F (motif B), or T/M/I/K/Y (motif C) at the P2 position, agreeing with the P2 specificity in the previous findings and the MEROPS database ^18^, except for T and K. Our motifs showed a clear preference for F at the P1′ position, which had not been reported before. The specificity motifs and the mass spectrometry results suggested that sequences containing consecutive hydrophobic amino acids (I, M, F and Y) are highly cleavable by streptopain, since each hydrophobic amino acid can be recognized as a P2 position resulting in multiple cleavage sites in a single sequence. A subsite cooperativity change was also observed among our motifs. In motif A, when Y was in the P1 position, M was preferred in P2 over other hydrophobic amino acids. In contrast, in motif B, when M was in the P1 position, F was preferred in P2 over other hydrophobic amino acids. A recent study reported the crucial cleavage of the GSDMA protein by streptopain after Q246 (ILIQ↓ASDV) which triggers pyroptosis ^56^. This cleavage after IQ was also supported by our data as demonstrated by high numbers of LC-MS/MS spectra for two peptides (Fig. 6b).

Since the producer of streptopain, *S. pyogenes*, causes a wide range of infections in human, we searched for matches of the 19 enriched sequences with MS-confirmed cleavage sites (Fig. 6b) in the human proteome database. Streptopain is known to target the proteins of the immune system and signaling proteins or systems of the host ^41^. The entire 6-mers and the 5-mer subsequences with confirmed cleavage sites, returned 3 and 230 matches, respectively. These matches included 37 cell surface proteins and 5 secreted proteins, such as complement C1r subcomponent (loop region), ion channels, receptors and glutathione peroxidase. Of course, despite being cleaved as short peptides, some of those matches might not be cleaved in the context of their native full-length protein and under physiologically relevant conditions. Therefore, such follow-up testing will be important but goes beyond the scope of the current study.

In summary, these results for streptopain vastly expand the previously published specificity information for this important protease. As with both streptopain and ADAM17, performing proteome-wide searches for sequences matching with more of our thousands of enriched peptide sequences, or against proteomes from other organisms, would likely generate additional hits for further studies.

In addition to vetting our method through confirming known substrate sequences as described above, we further validated our technique by demonstrating a strong correlation between relative enzymatic efficiencies previously reported and the high-throughput sequencing data output of our approach (Fig. S3). We compared the reported *k*_cat_/*K*_M_ values for ADAM17 substrates with highly diverse sequences (Fig. S3a), and substrates differing in only a single position (Fig. S3b) ^24^, with a substrate enrichment score calculated from the copy number of cleaved sequences in our high-throughput sequencing data. The same analysis was also performed for several streptopain substrates (Fig. S3c) ^42^. The strong correlations with Pearson correlation coefficients ρ ≥ 0.7 confirmed that the high-throughput sequencing data obtained with our method reflected the relative proteolytic kinetics of the substrates.

## Discussion and conclusions

The existing state-of-the-art methods have become adept at identifying canonical preferences of narrow-specificity proteases screens ^2-9, 12, 57-63^, yet, the cleavage preferences for broad-specificity proteases have been challenging to fully elucidate. The new specificity analysis approach described here has the following advantages over existing technologies: (1) We eliminated amino acid bias that usually results from using a proteomic substrate library isolated from an organism. Proteomic libraries do not present all possible permutations of amino acids ^2-6, 8, 32, 40^. Instead, we used a chemically synthesized library containing all possible octamer substrates rendering our method organism-independent and our results applicable to all organisms. The library was built from nucleic acid trimers instead of the more common degenerate NNS or NNK codons resulting in a more even amino acid distribution, free from detrimental stop codons. Analyzing the precise distribution of amino acids present in the library immediately before the protease cleavage allowed us to accurately compare enrichment of cleaved substrates. (2) Our library did not require protease processing prior to selection ^2, 4, 8, 32, 40^. PICS and some PICS-related methods require pre-processing with trypsin, for example, to prepare short peptide fragments leading to biases. Our library evenly presents essentially all possible octamers. (3) We included all natural amino acids without any modifications. Several previous methods omit amino acids like cysteine or methionine ^12^, or modify amino acids, such as protecting free lysine and cysteine residues ^2, 4, 8, 32, 40^. With mRNA display, we were able to assay all possible octamers as unmodified peptides. (4) By combining LC-MS/MS and high-throughput sequencing, we assessed both the prime and non-prime cleavage specificity simultaneously. We developed a computer script that incorporates the MS results into the alignment process and generates cleavage site-indicated motifs even without prior knowledge of the protease of interest. (5) We were able to test orders of magnitude more peptide substrates than most of the previous best methods ^32, 40^. (6) mRNA display is a purely *in vitro* technology, which, in principle, allows for modifications of cleavage conditions to easily assess changes in protease function with regards to temperature, pH, ionic strength, or inhibitors ^26^.

We applied this method to three proteases. The resulting detailed cleavage motifs were supported by testing more individual substrates than other current best methods. The specificity motifs for factor Xa markedly improved upon the previous best analysis by providing increased resolution at the P3 and P2′ sites ^32^. The specificity motifs generated for ADAM17 better resolved both sides of the cleavage position (Q at P1 and P1′ positions), compared to previous analyses ^40^. The new and detailed specificity motifs for streptopain, the previously least characterized of the three proteases tested here, will lead to a better understanding of this important protease than ever before ^42^. Besides the utility of the final specificity motifs, this method identified several thousand individual cleavable peptides for each protease that can be also used directly to search proteomes for sequence matches. Potentially interesting matches could be the starting point for future research that would need to include a confirmation that cleavage is also happening in the context of the whole native protein.

While our method overcomes a wide range of limitations, biases and shortcomings of previous approaches, there are some aspects that could be further improved. For example, potential impurities in the chemically synthesized peptides could affect the accuracy of the MS/MS cleavage site analysis. However, this potential issue can be minimized by either using high-purity peptides, or by increasing the minimum required number of spectra used for the MS/MS data-guided realignment. Furthermore, some octamer sequences in our comprehensive library may translate less efficiently than others and could therefore be underrepresented. We reduced this potential bias by avoiding rare codons in the library through trimer nucleotide synthesis ^30^ and by 30-fold oversampling of all possible octamers. However, some translation bias might persist for a few unusual octamer sequences. Similar to other cleavage specificity analysis methods, the detection of cleavage of a small peptide does indicate - but not guarantee - cleavage of the same sequence within the context of a larger, folded protein. Therefore, protease specificity analyses in general, including the method presented here, only yield likely predictions for protein cleavage that will ultimately have to be tested in native conditions.

In conclusion, the method described here yields specificity determinations with an unprecedented level of detail and can be applied to identify the specificity of potentially any protease. This advantage in throughput is particularly crucial to understand broad-specificity proteases. Since proteases are ubiquitous enzymes with critical roles in hormone activation, digestion, and immune response, and are the target of about 10% of all pharmaceuticals ^64^, this method has the potential to help address numerous medical problems, and provide a comprehensive analysis tool for protease engineering and potential inhibitor design studies.

## Author Contributions

M.D.L. performed mRNA-displayed peptide library design, preparation, and digestion; analysis of high-throughput sequencing data. B.Z. performed mRNA-displayed peptide library digestion; high-throughput sequencing samples preparation and analysis; subset peptide library design, preparation, digestion, and MS sample preparation; analysis of bioinformatics and MS data. J.A.B. developed and implemented bioinformatics approaches. T.J.G. helped conceive experiments for mass spectrometry and design the data analysis workflow; P.D.J. helped analyze mass spectrometry data and interpret results. M.D.L., B.S., and B.Z. designed the research, analyzed the data, and wrote the paper. All authors reviewed and edited the paper.

## Supporting Information

Supplementary tables, figures, and experimental details are available in the Supplementary Information.

## Acknowledgements

We thank M. F. Freeman, L. Higgins and T. W. Markowski for advice on mass spectrometry; Kun-Hwa Lee for optimizing the hammerhead cleavage protocol; A. J. O’Donoghue, P. M. Schlievert and B. Walcheck for scientific discussions; and J. D. Lipscomb for comments on the manuscript. This work was funded by grants from the US National Institute of Health (NIH) (AI113406; MSTP grant T32 GM008244 to M.D.L.; U24CA199347 to T.J.G. and P.D.J.), the Minnesota Medical Foundation (4036-9663-10 and 4088-9221-12), the University of Minnesota Biocatalysis Initiative, the Office of VP of Research at the University of Minnesota (Grant-in-Aid), University of Minnesota Informatics Institute (Updraft Grant), the American Heart Association (12PRE11900061 to M.D.L.), the Graduate School Doctoral Dissertation Fellowship (M.D.L.), the Center for Metabolomics and Proteomics at the University of Minnesota for generation of mass spectrometry data.

