## Supplementary Information for "Comprehensive protease specificity profiling"

### Contents

**Supplementary Tables**


1 Amino acid distribution of octamer library

2 Protease digest of octamer library at varying concentrations of protease

3 Number of reads from high-throughput DNA sequencing

4 Positions of motifs from Figs. 4, 5, and 6 within hierarchical trees

5 Nucleotide sequences of primers


**Supplementary Figures**


1 Detailed motif mining flowchart

2 Tree generation model

3 Correlations between substrate enrichment scores and protease catalytic efficiencies

4 Library design

5 tRNA purification gel

6 Peptide design for mass spectrometry


**Experimental Details**

**Supplementary Table 1.**

**Amino acid distribution of octamer library.**

The octamer library was chemically synthesized from Trimer Phosphoramidites and shows an amino acids distribution that is much closer to an even distribution of 5%, compared to libraries commonly synthesized from NNS or NNK degenerate codons. The color-coding is based on the mean centered values with darker shades representing a larger deviation from the desired mean (yellow). The octamer library from Trimer synthesis is free from stop codons, while about one quarter of NNS/NNK octamers contain a stop codon. The immobilized library represents the peptides immediately before the incubation with protease. The immobilized library is the result of transcription, translation and click-immobilization, which caused only minor changes in the amino acid distribution.

| Amino acids | NNS/NNK (theoretical) | Trimer synthesis (theoretical) | **Octamer library** Trimer synthesis (observed) | **Octamer library** Immobilized library (observed) |
|---|---|---|---|---|
| A | 6% | 5% | 4% | 4% |
| C | 3% | 5% | 5% | 5% |
| D | 3% | 5% | 5% | 4% |
| E | 3% | 5% | 5% | 4% |
| F | 3% | 5% | 6% | 7% |
| G | 6% | 5% | 3% | 3% |
| H | 3% | 5% | 5% | 5% |
| I | 3% | 5% | 7% | 7% |
| K | 3% | 5% | 5% | 5% |
| L | 10% | 5% | 4% | 5% |
| M | 3% | 5% | 5% | 4% |
| N | 3% | 5% | 4% | 4% |
| P | 6% | 5% | 5% | 5% |
| Q | 3% | 5% | 7% | 6% |
| R | 10% | 5% | 4% | 5% |
| S | 10% | 5% | 5% | 4% |
| T | 6% | 5% | 6% | 5% |
| V | 6% | 5% | 6% | 7% |
| W | 3% | 5% | 3% | 4% |
| Y | 3% | 5% | 7% | 8% |
| Stop | *3%* | 0% | 0% | 0% |

#### Supplementary Table 2.

**Protease digest of octamer library at varying concentrations of protease.**

| Protease | | Substrate-fusions / protease | Substrate-fusions displayed $\times 10^{11}$ | Background signal | Substrates cleaved |
|---|---|---|---|---|---|
| **Factor Xa** | Low | 72 | 9.4 | 0.11% | 0.20% |
| | Intermediate | 3.6 | | | 0.19% |
| | High | 0.18 | | | 0.21% |
| **ADAM17** | Low | 9.5 | 2.7 | 0.95% | 0.87% |
| | High | 0.48 | 2.7 | | 2.4% |
| **Streptopain** | Low | 480 | 8.6 | 0.03% | <0.05% |
| | Intermediate | 24 | | | 0.31% |
| | High | 1.2 | | | 3.7% |

**Supplementary Table 3.**

**Number of reads from high-throughput DNA sequencing.**

| **Sample** | | **Reads** | | |
|---|---|---|---|---|
| **Type** | **Source or protease concentration** | **Initial forward read** | **Processed** | **% Used** |
| Controls | Trimer synthesis library | 22,002,271 | 14,503,025 | 66% |
| | Immobilized library | 22,432,133 | 14,214,360 | 63% |
| Streptopain | Low 1/400x | 22,165,915 | 14,724,444 | 66% |
| | Intermediate 1/20x | 21,989,882 | 13,954,406 | 63% |
| | High 1x | 21,224,393 | 13,953,636 | 66% |
| Factor Xa | Low 1/400x | 21,740,644 | 13,446,391 | 62% |
| | Intermediate 1/20x | 27,425,979 | 16,604,651 | 61% |
| | High 1x | 22,167,802 | 14,290,899 | 64% |
| ADAM17 | Low 1/50x | 21,795,672 | 13,795,322 | 63% |
| | High 1x | 21,785,137 | 13,469,583 | 62% |

**Supplementary Table 4.**

**Positions of motifs from Figs. 4, 5, and 6 within hierarchical trees.**

| | Motif | Position |
|---|---|---|
| Fig. 4a | 1 | 01 |
| | 2 | 0001 |
| | 3 | 0010 |
| Fig. 4c | A | 00100 |
| | B | 00001 |
| Fig. 5a | 1 | 010001 |
| | 2 | 0011 |
| | 3 | 00101 |
| | 4 | 00100 |
| | 5 | 010000 |
| | 6 | 10100 |
| Fig. 5c | A | 00110011 |
| | B | 00100 |
| | C | 00110000 |
| Fig. 6a | 1 | 0100 |
| | 2 | 0000000 |
| | 3 | 1111 |
| Fig 6c | A | 00101 |
| | B | 001101 |
| | C | 0010010 |

**Supplementary Table 5.**

**Nucleotide sequences of primers.**

| Primer name | Sequence |
|---|---|
| D18 FW Ext 2 | 5′- TCTAATACGACTCACTATAGGGTTAACTTTAGTAAGGAGGACAGCTAAATG |
| 8mer RV Ext 4 | 5′- TTAATAGCCGGTGGGCATAGGCTGTG |
| 8mer RV Ext 5 | 5′- TTAATAGCCGGTGGGCATAG |
| Spec RT (PQPMP) V6 | 5′- TTTTTTTTTTTTTTTTCGGCATAGGCTGTGG |
| o-tRNA FW 1 | 5′- GCTTTTAGATCTTAATACGACTCACTATAGGG |
| pPaFT RV | 5′- TGGTCCGGCGGAGGGGATTTG |
| NGS FW V4 | 5′- CATCATCACCCGCAAC |
| NGS RV V4 | 5′- GCCGGTGGGCATAGG |

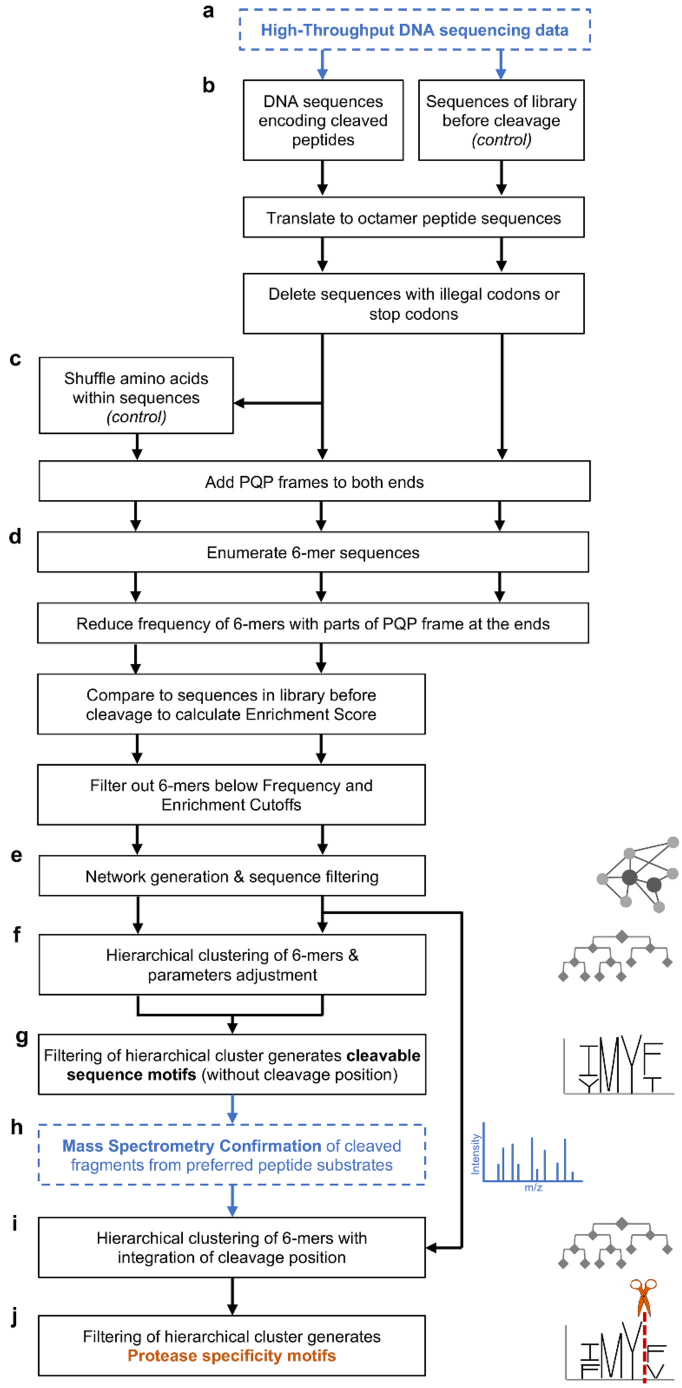


**Supplementary Figure 1. Detailed computational motif mining flowchart.**

Automated processing of the high-throughput DNA sequencing data from the peptide substrate library before and after protease cleavage, and the subsequent incorporation of mass spectrometry cleavage data for preferred peptide substrates ultimately yields the protease specificity motifs for a given protease. The two data entry points are highlighted by dashed blue boxes. The lettering **a** to **j** is identical to the less detailed overview Fig. 3.

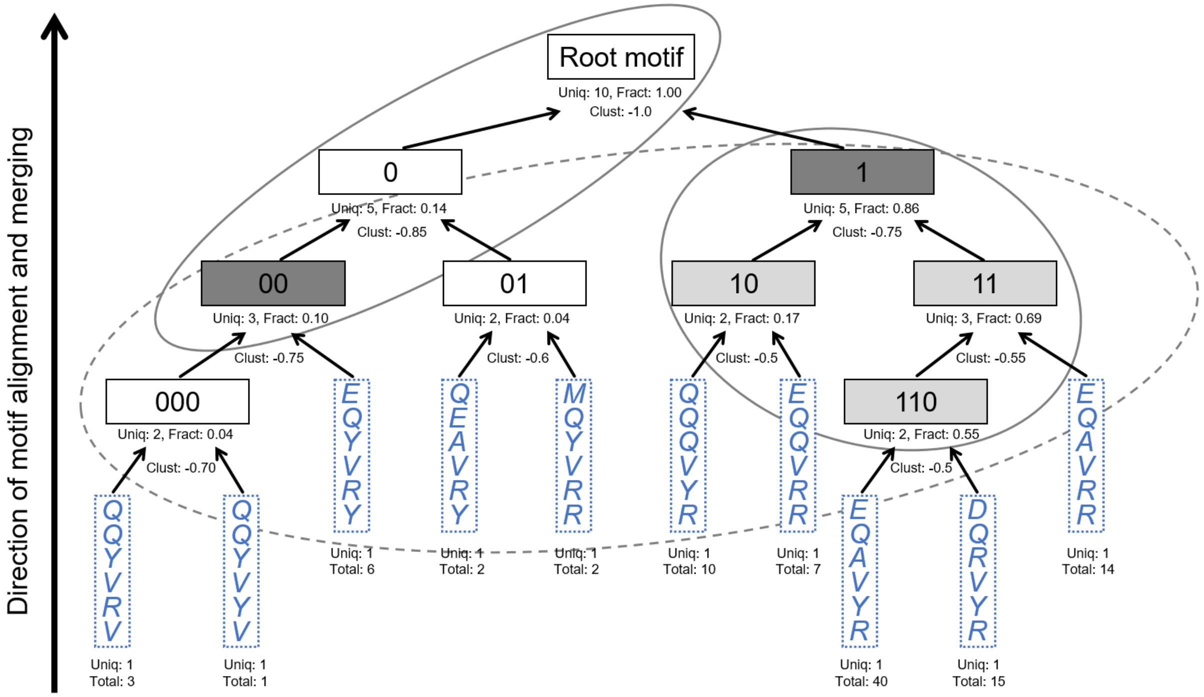


**Supplementary Figure 2.**

**A simplified model of motif tree generation and filtering.**

This figure only serves to visualize the general principles of the hierarchical tree generation and filtering approach. This is a made-up simplified tree as the actual trees generated for the three proteases are much more complex and too large to be displayed here. The input data for the motif tree generation were the processed peptide sequences obtained from DNA high-throughput sequencing shown in dashed blue boxes. Each black box represents a motif and the associated number in binary code shows the location of a motif in the hierarchical tree. Uniq: Number of unique 6-mer sequences supporting this motif. Total: Number of sequences supporting this motif. Fract: Sequences supporting this motif as fraction of all sequences enriched for this protease. Solid ellipses encircle all motifs supported by at least 5% of all enriched sequences (filtered for Fract > 0.05). Clust: Clustering metric representing similarity of the two motifs that were merged to yield this motif. A low clustering metric means that the motifs had low similarity. The dashed ellipse encircles motifs that were merged from two daughter motifs with certain minimal similarity (filtered for Clust > -0.8). After application of those two filters, the five motifs named 00, 1, 10, 11and 110 remained (highlighted in two shades of grey). Among motifs 1, 10, 11 and 110, the motif 1 was the parent motif and the motifs 10, 11 and 110 were daughters in a branch. Therefore, only the parent motif 1 was considered further as it also contained the daughter motifs information. In summary, only the motifs 00 and 1 would be shown as the motif mining result from this simulated model tree. The Clust metrics presented in the figure are mock numbers for illustration purposes only. In our study, the Fract and Clust filters were adjusted to yield 3-6 motifs for the subsequent processing in the motif-mining pipeline.

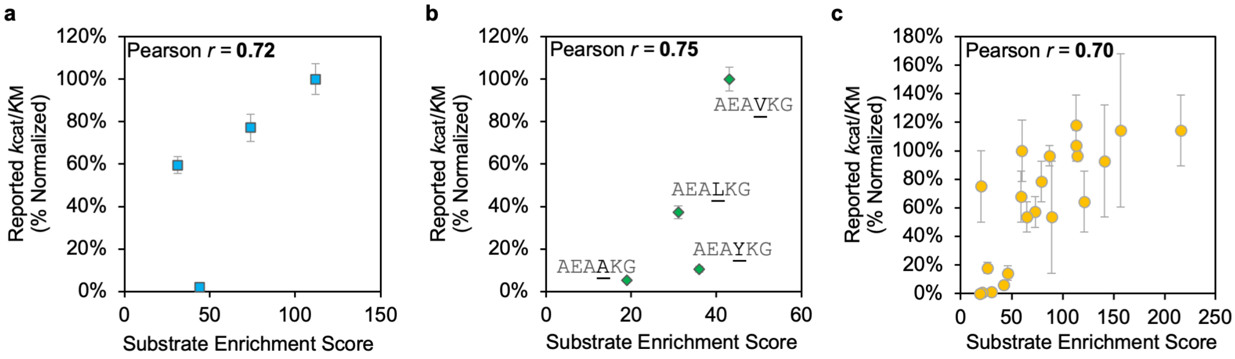


**Supplementary Figure 3.**

**Correlation of protease catalytic efficiencies with high-throughput sequencing data from the mRNA display workflow in this study.**

Catalytic efficiencies previously reported for ADAM17 for vastly different substrates (a) or substrates that differed only at the P1′ position (b) [1], as well as efficiencies reported for streptopain (c) [2], correlated well with the substrate enrichment cores determined in the current study. Pearson correlation coefficient *r* is shown in each panel, with p-value < 0.05.

| T7 promoter | TMV translation enhancer | Start | G | Amber Stop |
|---|---|---|---|---|
| 5′-TCTAATACGACTCACTATAGGG | TTAACTTTAGTAAGGAGGACAGCTAA | ATG | GGC | TAG |

| $His_6$ | PQP frame | | PQP frame | Linker | Stop |
|---|---|---|---|---|---|
| CATCACCACCATCATCAC | CCGCAACCG | $(Trimer)_8$ | CCACAGCCT | ATGCCCACCGGCTAT | TAA-3′ |
| | | Octamer library | | | |

**Supplementary Figure 4.**

**Nucleotide sequence of DNA library used to mRNA-display the comprehensive octamer peptide library.**

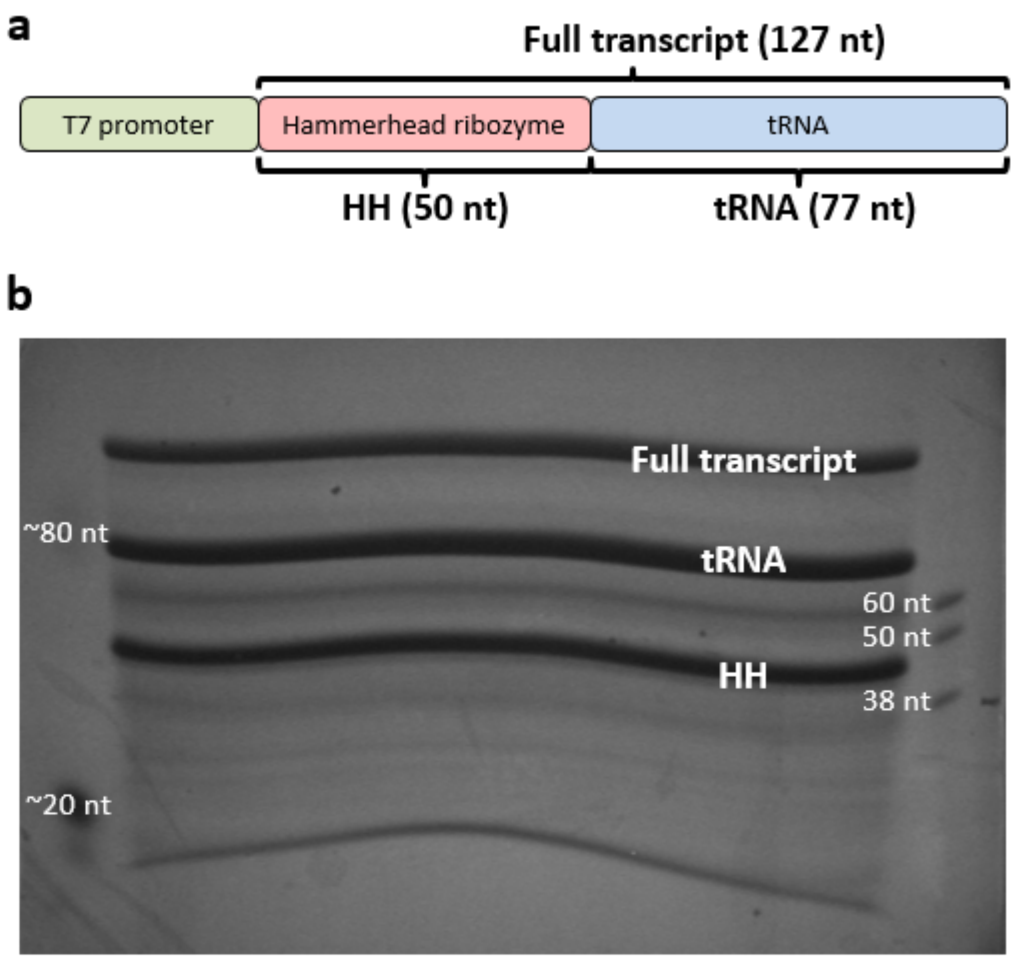


**Supplementary Figure 5.**

**Generation and purification of the pPaF-tRNA.**

**(a)** Design of the DNA template that was *in vitro* transcribed to RNA consisting of the hammerhead ribozyme and the pPAF-tRNA. **(b)** A denaturing polyacrylamide gel electrophoresis gel is shown that was used to separate the desired pPaF-tRNA (tRNA, 77 nt) from the uncleaved RNA transcript of the hammerhead ribozyme – pPaF-tRNA fusion construct (Full transcript, 127 nt) and the cleaved hammerhead ribozyme (HH, 50 nt).

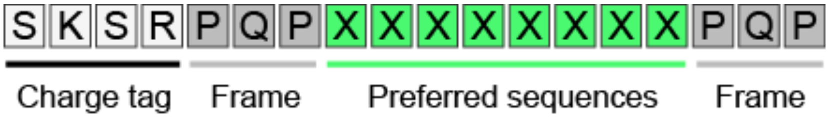


**Supplementary Figure 6.**

**Design of chemically synthesized peptides used for LC-MS/MS analysis of the cleavage position.**

Three types of charge tag peptides were fused to the N-terminus of the octamer substrates to add two (shown above), one or zero positively charged residues, SKSR, SSSR, and SSSS, respectively. The resulting 18-mer peptides were subjected to the protease digest and their cleaved fragments were subsequently analyzed by mass spectrometry.

#### Experimental Details

##### Chemicals.

All chemicals were purchased from Sigma-Aldrich unless otherwise stated.

##### Construction of comprehensive library of octamer peptides.

The octamer library (Fig. 2b, for full DNA sequence see Fig. S4) was synthesized by the Keck Biotechnology Resource Laboratory at Yale University using trimer phosphoramidites (Glen Research) for the random octamer region. The library was amplified with primers D18 FW Ext 2 and 8-mer RV Ext 5 (Table S5).

##### Expression and purification of p-propargyloxy-L-phenylalanine tRNA synthetase (pPaFRS).

We followed the published protocol for the cloning, expression, and purification of pPaFRS [3] with the following changes: (1) Cells were split into 4 × 250 ml aliquots, spun down for 15 min, and not washed prior to freezing at -80 °C. (2) A 250 ml aliquot was resuspended in lysis buffer. (3) Sonication was used to lyse the cells. (4) 1.5 ml of Ni-NTA resin was used in a gravity flow system. (5) 3.5K MWCO slide-a-lyzer (0.5-3 ml size) was used for dialysis. (6) The sample was dialyzed against 3 × 2 l of 1x PBS buffer for 2 h, 2 h, and overnight. (7) Recovered samples were filtered through a 0.45 μm Millipore Ultrafree PVDF centrifugal filter and not concentrated further.

##### Synthesis of pPaF-tRNA by transcription 5′-trimming using the hammerhead ribozyme, and its charging with p-propargyloxy-L-phenylalanine (pPaF).

The DNA encoding a tRNA sequence optimized for use with pPaFRS (o-tRNA$^{opt}$) and fused to a hammerhead ribozyme [4] was synthesized by GenScript and PCR-amplified with primers o-tRNA FW 1 and pPaFT RV (Table S5). The PCR product was agarose-gel purified and transcribed by T7 polymerase into RNA. To correctly trim the 5′-terminus of the tRNA using the hammerhead ribozyme, the RNA was mixed with $MgCl_2$ to 30 mM and Tris-HCl (pH 8.3) to 40 mM. This mixture was split into 100 μl aliquots, transferred to PCR tubes, and heat-cycled using the PCR Thermal Cycler program: (1) 1 min at 72 °C, 5 min at 65 °C, 5 min at 37 °C, repeat total of 15 times. (2) 2 min at 60 °C, hold at 4 °C. The product was ethanol precipitated and purified by denaturing 8% urea-PAGE. The dominant middle band (Fig. S5) was cut out, and the RNA was eluted from the gel by the crush and soak method to obtain purified pPaFT.

The pPaFT was charged with pPaF by combining the following: 100 mM HEPES-KOH (pH 7.2), 30 mM KCl, 12 mM $MgCl_2$, 4 mM ATP, 2 mM DTT, 1 mM pPaF purchased from Chem-Impex International Inc., 10 μM pPaFT, 3 μM pPaFRS. The reaction was split into 2.5 nmol pPaFT aliquots, incubated at 37 °C for 1 h, ethanol precipitated twice and stored dry at -80 °C until immediately prior to use.

##### mRNA display of octamer peptide library.

The transcription, cross-linking, and translation using rabbit reticulocyte lysate (Promega) were performed as described previously [5]. The translation using PURExpress® (NEB) and pre-charged tRNA was performed by combining the following for a 125 μl final reaction volume: (1) PURExpress manufacturer-provided "solution A Δ aa/tRNA", "solution B Δ RF123", tRNA, RF2, and RF3 according to the manufacturer's protocol. (2) Amino acid mixture Δ methionine from Promega to 100 μM. (3) $^{35}$S-labeled methionine to 0.344 μM. (4) Cross-linked RNA template to 1 μM. (5) Pre-charged pPaF-tRNA to 20 µM. The reaction was incubated at 37 °C for 2 h. KCl and $MgCl_2$ were added to final concentrations of 550 mM and 50 mM, respectively. The solution was incubated at room temperature for 30 min.

The mRNA-displayed peptides were oligo(dT)-purified and reverse transcribed as described previously [5], except that we used a 100-fold excess of reverse transcription primer Spec RT (PQPMP) V6, and 3,000 U/ml Superscript II. The mRNA-displayed peptide fusions were ethanol precipitated twice to eliminate the Tris buffer and to concentrate them.

**Immobilization of mRNA-displayed octamer peptide library by click-reaction with azide-agarose resin.**

The click immobilization strategy used here was derived from a published protocol [6]. Solution A was prepared in bulk and stored at 4 °C until use: 200 mM HEPES-KOH (pH 7.6), 10 mM aminoguanidine hemisulfate, 0.05% Triton X-100. Solution B was prepared fresh for every click immobilization: 2 mM $CuSO_4$, 2 mM Tris(3-hydroxypropyltriazolylmethyl)amine (THPTA). 350 µl of azide-agarose resin 50% slurry from ClickChemistryTools was rinsed with 3 × 1 ml $H_2O$, then blocked with 0.5 mg/ml yeast tRNA for at least 20 min. Four 0.5 ml tubes were cut to remove their caps. The resin was rinsed with 3 × 1 ml $H_2O$ then 3 × 1 ml of Solution B. 120 µl of Solution B was added to resuspend and transfer the resin to a capless 0.5 ml tube. 120 µl of Solution A was used to resuspend the ethanol precipitated peptide fusions and transfer them to a capless 0.5 mL tube. Another capless 0.5 ml tube was filled with 150 µl $H_2O$. 1.5-2 mg of sodium L-ascorbate was weighed into a capless 0.5 ml tube. The two tubes containing either $H_2O$ or ascorbate were placed in a 25 ml heart-shaped flask, while the tubes containing Solution A including peptide fusions or Solution B were placed in a 50 ml heart-shaped flask. The flasks were purged under argon flow for 1 h. 50 µl $H_2O$ was added to each 1 mg of ascorbate in the purged flasks and suspended. 24 µl of the suspended ascorbate was transferred to the Solution B tube. All of Solution A was transferred to the Solution B tube. The reaction was incubated at room temperature for 3 h while resuspending the resin once per h.

**Washing of immobilized octamer peptide library.**

The resin slurry from the click reaction was recovered and transferred to a 10 ml Poly-Prep chromatography column (Bio-Rad). The resin was carefully washed with 8 × 500 µl, then 4 × 10 ml of 60 °C warmed denaturing urea buffer (8 M urea, 500 mM NaCl, 20 mM Na-Phosphate (pH 6.5), 0.1% Triton X-100). Each wash was measured by scintillation counting to track the $^{35}S$-labeled peptides.

The resin was washed with the appropriate digestion buffer for the respective protease that it would be digested with subsequently: (1) Factor Xa: 50 mM HEPES-KOH (pH 7.4), 150 mM NaCl, 1 mM $CaCl_2$, 0.1% Triton X-100. (2) ADAM17: 50 mM HEPES-KOH (pH 7.4), 2.5 µM $ZnCl_2$, 0.1% Triton X-100 and additionally containing 150 mM NaCl. (3) Streptopain: 50 mM HEPES-KOH (pH 7.4), 150 mM NaCl, 0.1 mM EDTA, 0.1 mM DTT, 0.1% Triton X-100. Note that ADAM17 digestion buffer usually does not contain NaCl, but this addition was necessary to achieve sufficiently high ionic strength for efficient washing. 10 × 1 ml, then 4 × 500 µl of digestion-wash buffer was used, while scintillation tracking decrease in signal. Next, the column was capped and 800 µl (factor Xa, streptopain) or 600 µl (ADAM17) of digestion buffer was added. The column was rotated at 37 °C for 1 h, then uncapped and flow-through collected. Each column was washed for 2-4 times with 500-800 µl of digestion buffer. This column wash was repeated, and all fractions were scintillation-counted until the wash fraction contained 0.1% or less of immobilized signal. The number of washes needed to reach this goal varied significantly between the three digestion buffers. Streptopain buffer (containing 0.1 mM EDTA and 0.1 mM DTT) reached 0.03% background after a single overnight wash and 3 subsequent 1 h washes. In contrast, factor Xa buffer (containing 1 mM $CaCl_2$) required two overnight washes and 11 1 h washes to reach 0.12% background. The radiation counts measurement from the final 1 h wash was used to define the background signal for the subsequent protease digestion.

**Protease digestion of immobilized octamer peptide library.**

We produced purified and highly active streptopain through our previously published method [7]. For both factor Xa (New England Biolabs) and streptopain, the digestions were performed at three concentrations of protease to achieve cleaved peptide signal above background (Table S2). 800 µl of the respective protease digestion buffer was added to the immobilized and washed peptide library and then the lowest amount of protease was added. The column was capped and rotated at 37 °C, then uncapped and the flow through containing cleaved fusions was collected. This process was repeated for the intermediate and high protease concentrations. Protease concentrations were initially based on reported manufacturer recommendations or published activities and then modified to achieve a range of cleaved signals below and above background signal. For streptopain, 82 pg, 1.6 ng, and 33 ng were added for low, middle, and high digestions. For factor Xa, 250 pg, 5 ng, and 100 ng were added for low, middle, and high digestions. The factor Xa digest resulted in weakly increased signals above background at all protease concentrations. Streptopain digestions released cDNA-fusions above the background signal at the intermediate protease concentration. The elutions of cleaved peptide-mRNA-cDNA fusion molecules were purified and concentrated with a QIAquick PCR purification kit (Qiagen). The cDNA of the fusions was amplified using NGS primers NGS FW V4 and NGS RV V4 (Table S5).

For digestions with ADAM17 (R&D Systems), the washed peptide library resin was divided into three equal parts and each of those aliquots was transferred into a separate column. This separation was necessary because the ADAM17 digestion must not contain NaCl, which inhibits digestion. The resin in each column was washed with digestion buffer (50 mM HEPES-KOH (pH 7.4), 2.5 µM $ZnCl_2$, 0.1% Triton X-100), then resuspended in 200 µl of digestion buffer. To one each of the three columns, either buffer-only, low ADAM17 protease (1 ng), or high ADAM17 protease (50 ng) was added. The digestion was incubated with rotation for 16 h at 37 °C. Any cleaved peptide-mRNA-cDNA fusion molecules were eluted from the agarose resin with 600 µl of elute buffer (50 mM HEPES-KOH (pH 7.4), 200 mM NaCl, 2.5 µM $ZnCl_2$, 0.1% Triton X-100). The ADAM17 digestion was above background only at the highest protease concentration tested. The cleaved peptide-mRNA-cDNA fusions were purified, concentrated, and PCR-amplified as described for the other two proteases in the paragraph above.

**High-throughput sequencing of cDNAs that encode cleaved octamer peptides and raw read processing.**

The immobilization library was first amplified using primers D18 FW Ext 2 and 8mer RV Ext 4 from resin before amplifying with NGS primers NGS FW V4 and NGS RV V4 (Table S5). The starting library, the immobilization library and the eluted encoding cDNA after digestion were sequenced by ACGT, Inc. using an Illumina NextSeq500 with a paired end, 75 bp read. Approximately 21-27 million DNA sequence reads each for the library of peptides before and after protease cleavage were obtained by high-throughput sequencing. The data were processed with the following programs using the Galaxy server: (1) PEAR alignment to match forward and reverse reads. (2) Manipulate FASTQ to reverse-complement incorrectly oriented sequences. (3) Manipulate FASTQ to require 100% identity on the 5′ and 3′ flanking regions of the octamer, and trim off these nucleotides. (4) Filter FASTQ all sequences that are not exactly 24 nucleotides. The data resulting from step (4) (Table S3) were used as input for the motif mining program described below. For the following analyses we used the datasets obtained from the protease digests at the lowest concentration of protease that yielded > 2-fold cleaved substrates over background.

**Computational motif mining.**

The DNA (24 bp) encoding the octamer peptide library had been chemically synthesized from nucleotide trimers using only a single codon per amino acid. As a result, the presence of either an improper codon or a stop codon in the sequencing data indicated a strong likelihood of sequencing error, or sequence misassembly. All DNA sequences containing only correct codons were translated into their octamer peptide sequences (8-mers).

A detailed schematic overview of the entire computational motif mining is shown in Fig. S1. In addition to the octamer dataset obtained from sequencing, a control dataset was generated *in silico* by shuffling the amino acid order of the sequences in the selected pool. The control was later used to estimate the false positive rate of the detection method. The 8-mer amino acid sequences resulting from steps a and b represented only the variable region of the cleavable peptide. In addition, there were three constant amino acids, ‘PQP’, on each end framing the variable region during the protease digestion. For the bioinformatics analysis, the PQP frames were added back to both ends of the octamers to regenerate the 14-mer peptide sequences as they had been displayed in the cleavage experiment. This was necessary to account for the unlikely event that a protease partially recognized, or cleaved within, the PQP frame sequence. A k-mer size of 6 was used as the optimal length for the primary analysis and the k-mer frequencies were tallied. Longer k-mer sizes did not allow for a sufficient dynamic range in the k-mer frequencies. The use of shorter k-mers would have reduced the ability to detect coordination between distant positions. Due to an expected overrepresentation of 6-mers beginning with ‘P’ or ‘QP’, or ending with ‘P’ or ‘PQ’, an additional correction was added. 6-mers that overlapped by one amino acid with the static frame region were treated as 1/20th (20^1) of a k-mer, those that overlapped by 2 amino acids were treated as 1/400th (20^2), and those that overlapped by 3 as 1/8,000th (20^3). 6-mers from protease-treated samples were filtered from further analysis based on a frequency threshold and their enrichment score relative to the immobilized library control (peptides before cleavage). For the enrichment score calculation, a pseudo count of 1 was added to all k-mers to account for sequences that were found in the enriched cleaved library but not identified in the much more complex library before cleavage. This was necessary because the complexity of this starting library far exceeded the sequencing depth of high-throughput sequencing.

The minimum detection frequency for k-mers was set on a dataset-by-dataset basis and was largely influenced by the depth of sequencing. This value was selected to retain a similar number of k-mers between the samples. It ranged from 6-14.

Similarly, minimum enrichment was set to retain a consistent number of k-mers to progress to the network processing stage. This number was also dependent on sequencing depths. Increased retention of k-mers is expected to increase sensitivity of motif detection. However, the pairwise comparison of the k-mers used to generate the network is demanding both in terms of computational time and memory capacity. In order to enable generations of large numbers of networks quickly we required 6-fold enrichment in most cases, though for some datasets increased this to as high as 12-fold. The number of unique 6-mers retained at this stage of the workflow for factor Xa, ADAM17 and streptopain was 2,693, 4,861 and 2,954, respectively.

Based on the hypothesis that protease targeting will not be defined by a single fixed 6-mer peptide, but instead by a subsequence with a degree of variability, we sought to enrich the k-mer set for k-mers with similar sequences. To do so, a network was created with retained k-mers as nodes and weighted edges based on the pairwise distance between nodes. The edges were calculated as the ungapped alignment distance between the peptide sequences. While multiple measures of similarity were evaluated, ultimately a simple ungapped alignment was used with a similarity matrix scoring matching amino acids as +2 and mismatching amino acids as -2. Also, amino acid matches to end gaps were scored as a -2. Additional matrices are possible where amino acids are grouped based on similarity of characteristics. Here +1 is used for a within group match. The resultant alignment score is converted to a similarity score with a value of 1 indicating identical strings and a value of 0 indicating no shared positions.

Numerous cutoffs for node degree and edge weight were tested to identify values that consistently gave large complex networks with the true datasets but gave sparse or no networks with randomized datasets. We found that a degree cutoff of 4 and an edge weight cutoff of 0.55 gave consistent results. After applying these filters, 2,369, 1,059 and 2,015 unique 6-mers were retained for factor Xa, ADAM17 and streptopain, respectively.

In order to determine consensus motifs from the enriched set of k-mers, the remaining k-mers were hierarchically clustered using a CLINK-like approach [8]. In this case, the k-mers were linked agglomeratively into position weight matrices (PWMs) based on a metric where the ungapped alignment was computed for each pair of k-mers between the two PWMs. The metric used for scoring the alignments was the same as was used for the earlier network generation. This metric averaged across all pairs and weighted by the observation frequency of each k-mer. The same workflow was run on the shuffled controls. In most cases, these runs ended at the network generation stage. Given the degree requirement, without an enrichment of some similar k-mers, no network is retained. In a minority of cases, some sort of network survived. In that case, a motif was generated at the end. These were visually compared to the true datasets to ensure that observed motifs were not present in the random data.

Selection of clusters based on similarity and the proportion of the total data represented allowed for the identification of motifs that represented subsets of the total cleaved sequence diversity (Fig. S2). To identify a subset of unique motifs, we filtered the motif sets based on the following criteria. First, all motifs had to be made up of at least 5% of the total unique k-mer population. This fraction filter was also used to compare the unique sequence count between samples. Secondly, motifs were only retained if they were formed from a merge that had a clustering metric (a measure for similarity) greater than -0.8, or -1.0 (streptopain dataset after including cleavage information), and were not also part of a motif with a greater proportion of k-mers that also met the clustering metric criteria. These motifs were visualized as octamers using WebLogo3 [9] (Figs. 4, 5, 6). The k-mers backing the generation of these motifs were used as guides for generating a sublibrary of peptides that were digested again and evaluated by MS/MS to determine their cleavage sites.

**Choosing a subset of enriched octamer peptide substrates for cleavage analysis by mass spectrometry.**

To determine the cleavage position of the highly enriched cleavable octamer sequences, we chemically synthesized about 1-2 dozen peptides for each protease, digested them with the respective enzyme, and analyzed the cleaved fragments by mass spectrometry. For each protease, we chose the five most enriched 6-mer sequences and the five most enriched 6-mer sequences from each cleavable sequence motif (panel a in Figs. 4, 5, 6). This was necessary because in rare cases, the top five sequences from the top cleavable motifs did not comprise all of the overall most enriched sequences. Nevertheless, any duplicate sequences among them were eliminated. We also included the two least enriched 6-mer sequences as controls. The confirmation of cleavage for those least enriched sequences implied that all of the even higher enriched sequences were cleavable too. We then chose the original octamer sequences that contained these 6-mer sequences from the high-throughput DNA sequencing data using an octamer sequence picking script. This script automatically applied the following four criteria. 1) the 6-mer sequence is located in the center of the octamer sequence; 2) of those octamers, any repeated sequence is chosen without further testing; 3) if there is no repeated octamer sequence or there are more than one repeated octamer sequence, the next criterion is applied; 4) the hydropathy score [10] of the remaining octamer sequences is calculated, and the sequence with the lowest score (highest solubility) is chosen. Next, we added back the PQP frame to the N- and C-terminus of each octamer to produce a 14-mer sequence.

In order to enable facile detection of the N-terminal cleavage fragments by mass spectrometry, we applied the following steps to ensure that the substrate peptides and the resulting cleaved fragments have a mass that is suitable for detection, carry a proper positive charge, and will likely be soluble. We built these steps into an Excel script for ease of execution. First, we calculated the maximum charge of the 14-mer sequences at pH 1 by summing up the number of K, R and H residues and adding 1. Depending on whether the calculated maximum charge was greater than 3, exactly 3 or less than 3, we added sequences SSSS, SSSR or SKSR to the N-terminus of each 14-mer, respectively (Fig. S6). These additions also increased the solubility of the peptide. The resulting 18-mer sequences were then examined for meeting the following three criteria. Criterion 1 requires the lowest m/z ratio of the 18-mer sequence to be between 380 and 1,800. Criterion 2 stipulates that the maximum charge of the N-terminal 11-mer fragment (assuming cleavage in middle of octamer region) must be greater than 1. Criterion 3 requires the lowest m/z ratio of the N-terminal 11-mer sequence to be between

380 and 1,800. Failure to meet any one of those criteria indicated the need to adjust the properties of the peptide by adding a different N-terminal 4-mer tag.

**Protease digestion of peptide sublibraries containing select preferred substrates.**

The 18-mer substrate peptides were synthesized by Genscript or Peptide 2.0 and obtained as crude peptides without further purification. As a positive control, a peptide was included that had previously been confirmed as substrate (for ADAM17: TNF-alpha PLAQA↓VRSSS [11]; for streptopain: SpeB autoprocessing site AAIK↓AGAR. The lyophilized peptides were re-suspended in deionized water to a final concentration of 5 mM. Crude peptides were mixed in equimolar ratios to create the sublibraries of up to 20 peptides (factor Xa: 17 preferred peptides; ADAM17: Two groups of 16 preferred peptides + 1 previously known substrate; streptopain: 19 preferred peptides + 1 previously known substrate). The peptide mixtures were desalted using C18 STAGE-tips as previously described [12] with the following modifications: an 18-gauge blunt-ended syringe needle was used to punch the core from C18 reversed-phase extraction disks (3M), and two cores were pressed into each syringe needle for extra loading capacity. The STAGE-tip was equilibrated with 20 µl of solvent 1 (80:20:0.1, acetonitrile:water:trifluoroacetic acid) and rinsed by 60 µl of HPLC grade water before loading the sample. 40 µg (~20 nmol) of peptide mixture was loaded into the STAGE-tip and washed twice with 60 µl of HPLC grade water. The peptides were eluted with ~40 µl of solvent 2 (60:40, acetonitrile:water). 16 nmol of purified peptides were aliquoted and dried in a SpeedVac concentrator.

The peptide sublibrary for factor Xa was re-suspended in 80 µl of FXa cleavage buffer (50 mM HEPES-KOH [pH 7.4], 150 mM NaCl, 1 mM $CaCl_2$), and 20 µl (4 nmol) of sublibrary was incubated with 3.48 µl of factor Xa (NEB, P8010, 1 mg/ml, 23 µM) at 37 °C for 1 h (protease:peptides, 1:50). The peptide sublibrary for ADAM17 was re-suspended in 1.6 ml of A17 cleavage buffer (20 mM Tris-HCl [pH 7.4], 2.5 µM $ZnCl_2$), and 19.2 µl (192 pmol) of sublibrary was incubated with 5 µl of ADAM17 (R&D Systems, 930-ADB, 0.2 mg/ml, 3.84 µM) at 37 °C for 15 h (protease:peptides, 1:10). The peptide sublibrary for streptopain was re-suspended in 80 µl of streptopain cleavage buffer (20 mM Tris-HCl [pH 7.4], 150 mM NaCl, 0.1 mM EDTA, 0.1 mM DTT), and 20 µl (4 nmol) of sublibrary was incubated with 2.74 µl of streptopain[7] (0.8 mg/ml, 29.2 µM) at 37 °C for 1 h (protease:peptides, 1:50). To prepare a no-protease negative control sample, a second aliquot of each peptide sublibrary was incubated under the same conditions in buffer only. The digestion reactions were stopped by the purification step described in the next section.

**Mass spectrometry (LC-MS/MS) of cleaved sublibraries to determine cleavage positions.**

To remove the polymer contamination of the commercial ADAM17 protease, its digest reactions were purified using a strong cation exchange (SCX) Zip-Tip (EMD Millipore) following the detergent removal protocol in the product manual except for adding 2% TFA solution to the sample before the clean-up. The elution was re-suspended in 20 µl of HPLC grade water. The cation-exchange purified ADAM17 sample and the crude factor Xa and streptopain digests were then purified by C18 STAGE-tip as described above, using 20 µl of HPLC grade water to rinse the reaction tube and load the sample onto the STAGE-tip. The peptides were eluted with 20 µl of solvent 2 and dried in a SpeedVac concentrator.

We analyzed the digested and purified peptide sublibraries on a capillary liquid chromatography (LC) Velos Orbitrap mass spectrometer system (ThermoFisher Scientific, Waltham, MA) as previously described [13] with the following adjustments: We injected approximately 2 pmol of each peptide in a volume between 2 and 5 µl. The LC column had an inner diameter was 100 µm and a length of 14 cm. The elution was performed at a flow rate of 330 nl/min with a mixture of solvent A (98:2:0.01, water: acetonitrile: formic acid) and solvent B (98:2:0.01, acetonitrile: water: formic acid) using the following gradient for solvent B: 2-8% over 2 min, 8-35% over 40 min, 35-90% over 1 min and 90% for 15 min. The mass spectra were acquired with the following spectrometer settings: no lock mass employed, MS1 range 380 – 1,800 m/z ratio, MS1 maximum injection time 300 ms, dynamic exclusion maximum number of values 300, duration 30 s and exclusion mass tolerance ± 15 ppm.

**Analysis of LC-MS/MS data to identify peptide fragments.**

The raw data acquired by LC-MS/MS were converted to mgf format via MSConvertGUI (ProteoWizard 3.0.11148 [14]). The peptide sequence databases were generated by combining the sequences of the cleaved sublibraries with the decoy sequences, which were the reversed sequences of the sublibraries. SearchGUI 3.3.20 [15, 16] was used for database searching with X!Tandem search engine [17] and default parameters except for the following changes: For the spectrum matching parameters, the unspecific digestion option was selected, and precursor and fragment *m/z* tolerant were set to 10 ppm and 0.02 Da, respectively. The precursor charge range was set from 2 to 6. The variable modifications setting for factor Xa and streptopain included oxidation of Met, Lys and Pro, acetylation of Lys, and deamidation of Gln and Asn; and the setting for ADAM17 included methylation of Lys, Arg and Ser, and dimethylation of the N-terminus. The import filter in the advanced setting allowed only 8-30 amino acid peptides in the search process. The false discovery rate (FDR) limits of the peptide and peptide spectrum match (PSM) were set to 1% in the validation level setting. Peptideshaker 1.16.12 [18] was used for post-processing with default settings. The chemical synthesis of peptides proceeds from C-terminus to N-terminus and regularly also yields truncated peptides as side products. To filter out undesired mass signals from truncated synthesis impurities, only the validated PSM numbers of the N-terminal fragment of the 18-mer peptide after cleavage were plotted against the position to show the cleavage position information. The PSM was treated as validated when 1) the identified peptide was not ‘invalid’; and 2) the PSM was either ‘confident’ or ‘doubtful’ with a confidence score of 100; and 3) all modifications were ‘confident’. If any MS spectra were detected in the negative control experiment without protease, the number of those spectra for a given fragment was subtracted from the number of the respective spectra for the protease digest. After the subtraction, if the largest number of spectra for any N-terminal fragment detected in the protease digest is equal or smaller than the absolute value of the number of spectra for any N-terminal fragment from the negative control, the MS results for this peptide were abandoned. From here on, the N-terminal peptide fragments, for which this subtraction resulted in ≤0 were not further considered. The user can optionally increase the stringency at this step to only focus on the major cleavage sites and motifs. The MS spectra results were exported to the following bioinformatics process.

**Protease specificity motif refinement using cleavage position information.**

Using cleavage site information from the MS/MS experiments, the afore described hierarchically clustering was rerun with the cleavage sites provided where a k-mer matched perfectly to a tested peptide. These cleavage sites were propagated up the tree and took precedence over the optimal score when determining how to align two sequences. In cases where multiple cleavage sites occurred in the same peptide, or a k-mer matched to multiple cleaved peptides, the occurrence rate of the k-mer was divided proportionally to the cleavage rate across all of the cleavage sites. Besides the fraction and similarity criteria described in the motif mining method, we chose to focus only on motifs that were supported by at least 10% of the total number of spectra from a sublibrary digestion to ensure substantial support for motif generation (FXa = 25;  A17 = 10; SpeB = 36).

**Calculation of substrate enrichment score.**

The substrate enrichment score used for the correlation with previously reported enzymatic efficiencies for a variety of protease substrates was derived from the cleaved octamer sequences. To cover over 99% of substrate sequence, a list of 4-mer sequences contained within the cleaved octamers was generated through sliding-window extraction. The number that a given 4-mer was found in all cleaved octamers was defined as the substrate count of that 4-mer. The substrate enrichment score was calculated by summing the substrate counts of three 4-mers derived from the P3–P3' region of a substrate through sliding-window extraction.
